# Moderate-Intensity Intermittent Training Inhibits miR-34a-5p to Reduce Prefrontal Cortex Apoptosis and Improve Cognitive Ability in Aging Rats Reduced by D-galactose

**DOI:** 10.1101/2025.08.19.671180

**Authors:** Qiaojing Gao, Jinmei Zhang, Liting Lv, Jun Gao, Meng Li, Xue Li, Lu Wang

## Abstract

Changes in microRNA (miRNA) play a role in brain aging. They are considered potential therapeutic targets. Regular long-term exercise benefits brain health. However, its exact mechanism is not fully understood. This study explored how moderate-intensity intermittent training (MIIT) improves cognitive function and reduces apoptosis in the aging brain. This study induced aging in rats by giving them D-gal injections (150 mg/kg/day) for 6 weeks. After confirming the model, the rats underwent MIIT. They exercised 45 minutes a day, 5 days a week, for 8 weeks. The results from behavioral, morphological, and molecular tests showed that 8 weeks of MIIT significantly slowed the decline in spatial learning and memory in D-gal aging rats. The exercise also improved the structure of the prefrontal cortex (PFC) and reduced apoptosis. Aging caused an overexpression of miR-34a-5p in the prefrontal cortex. MIIT reduced this overexpression. It also up-regulated Notch1 and inhibited excessive apoptosis by regulating Bcl-2 and Bax expression. In conclusion, MIIT may improve brain health by targeting miR-34a-5p and regulating apoptosis-related pathways.

## Introduction

Aging is a long and complex process. It happens when, as people grow older, tissues and organs gradually lose their function and shape because of cell aging. According to the World Health Organization, the population of people aged 60 and above is expected to double to 2.1 billion by 2050. By then, those aged 80 and over will number 426 million^[1–3]^. Today, thanks to advances in medical science and technology, people live much longer. Unfortunately, many suffer from chronic diseases. These conditions lower the quality of life in later years. Therefore, attaining healthy aging has emerged as a significant challenge for contemporary society. Among various chronic conditions, neurodegenerative diseases are particularly prominent, with brain aging constituting a key factor in their development^[4]^.Aging of brain regions such as the hippocampus, prefrontal cortex, and striatum is associated with impairments in cognitive functions, learning, and memory^[5,6]^. Although current research predominantly emphasizes the hippocampus, the prefrontal cortex—which is integral to memory, emotion, cognition, and decision-making—also warrants further investigation^[7–9]^.

Many studies have established that exercise is a promising nonpharmacological intervention with minimal side effects. Concurrently, the beneficial effects of physical activity on cognitive function have garnered significant attention^[10–12]^. Our earlier research demonstrated that moderate-intensity intermittent training enhances the synaptic structure and functional plasticity of the hippocampus in aging rats by activating the cAMP/PKA/CREB signaling pathway and up-regulating brain-derived neurotrophic factor expression, thereby improving spatial learning and memory[13]. Exercise appears to mitigate cognitive decline in both humans and animal models, reducing age-related memory loss and delaying the progression of neurodegenerative diseases. However, the precise mechanisms underlying the cognitive benefits of exercise remain to be elucidated. In recent years, research has identified microRNAs (miRNAs) as key mediators in processes related to physical exercise^[14]^. miRNAs are modulated by environmental factors and external stimuli, such as exercise, and they subsequently induce regulatory changes in transcriptional activity that impact a range of physiological and pathological processes, including cancer and aging^[15,16]^. For example, in the aging brain, miRNAs alter gene expression patterns to achieve neuroprotective effects that mitigate the aging process^[17,18]^. As a result, miRNAs have become a focal point in studies exploring the mechanistic underpinnings of aging-related chronic diseases^[19,20]^.

miRNA comprises small non-coding RNA molecules, typically 18-25 nucleotides in length, that regulate gene expression. These molecules inhibit mRNA translation by binding to the 3′ untranslated regions of target mRNAs, playing a central role in gene regulation at the transcriptional level^[21]^. Notably, miR-34a is significantly associated with aging; its elevated expression in aging brain tissue suggests that an increase in miR-34a may be a key factor contributing to the aging process^[22]^. It has been reported that miR-34a-5p plays a crucial role in various cellular processes and signaling pathways involved in multiple diseases^[23,24]^. Notably, miR-34a-5p modulates the Notch signaling pathway, thereby influencing cell proliferation and apoptosis, which in turn contributes to disease onset and progression^[25]^. Furthermore, activation of Notch1 has been shown to effectively inhibit neuronal apoptosis^[26]^. In studies involving SAMP8 mice, an elevation in hippocampal miR-132 levels was observed in untreated animals; however, exercise intervention downregulated miR-132 expression and reduced amyloid precursor protein (APP) accumulation, thereby mitigating hippocampal degeneration and enhancing cognitive function^[27]^. Similarly, chronic treadmill exercise increased cerebral cortical miR-124 expression, which corresponded with a reduction in amyloid-β (Aβ) accumulation^[28]^.

Therefore, whether exercise can activate Notch1 by down-regulating miR-34a-5p, reduce the excessive apoptosis of prefrontal cells induced by D-galactose (D-gal) in aging rats, and improve their cognitive and learning and memory ability has not been reported in the literature. Therefore, this study attempts to explore the effects and possible mechanisms of moderate-intensity intermittent training (MIIT) on miR-34a-5p/Notch1 and apoptosis in the prefrontal cortex (PFC) of D-gal-induced aging model rats, in order to provide effective therapeutic target screening and experimental basis for exercise to delay aging-related diseases.

## Materials and Methods

### Animals

Three-month-old male Sprague Dawley (SD) rats were purchased from Chengdu Dashuo Animal Experiment Co., Ltd. (Chengdu, China). Each rat weighed about 300±20g. They were kept at a temperature between 23-25°C with a humidity of 40-60%. The animals lived in a well-ventilated room with regular day and night cycles. They had free access to food and water. The experimental protocol received approval from the Academic Committee of Chengdu Sport University (Approval No. 2021-3).

After one week of adaptive feeding, A total of 36 rats were randomly divided into three groups: control (C), D-galactose (D), and D-galactose+exercise (DE). Each group had 12 rats. Fig 1 shows the entire experimental process.

**Fig 1.**
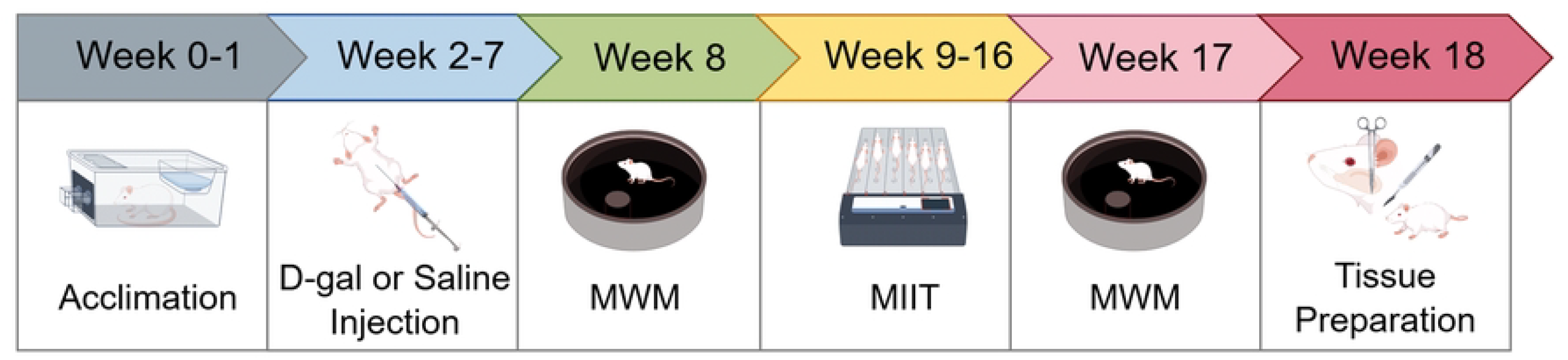
Protocol for the animal experiment. Week 1 was used for adaptive feeding. From weeks 2 to 7, subjects received daily intraperitoneal injections of either D-gal (at 150 mg/kg/day) or normal saline, totaling 6 weeks. In week 8, they spent one day adapting to swimming before the water maze experiment. The formal test included 6 days of positioning cruise experiments and 1 day of space exploration tests. Between weeks 9 and 16, an exercise intervention was carried out. In week 17, the water maze experiment was performed. After all tests were complete, subjects were anesthetized and tissues were collected.

### D-gal-Induced Aging Rat Model

D-galactose(D-gal) (IG0540) was purchased from Beijing Sunshine Biotechnology Co., Ltd. (Beijing, China). According to previous studies^[29]^, D-gal was dissolved in normal saline (0.9% NaCl). It was administered by intraperitoneal injection. The D-gal and D-gal + Exe groups received 150 mg/kg per day for 6 weeks. Meanwhile, the Ctrl group was injected with the same volume of normal saline.

### Exercise Protocols

The MIIT was adapted from the Bedford sports model^[30]^. The animal treadmill (SA101) used in this experiment was purchased from Cylons Biotechnology Co., Ltd. (Jiangsu, China). First, the rats in the DE group undergo one week of adaptive training on a treadmill. They run for 10 minutes each day at a speed of 10 m/min. Next, the formal intervention starts. The rats complete eight moderate-intensity interval exercise sessions per week for five days. The program begins with a warm-up at 10 m/min for 10 minutes. Next, the rats run at 20 m/min for 4 minutes and then slow down to 12 m/min for 3 minutes. They alternate these speeds for five rounds. In total, each treadmill session lasts 45 minutes with a slope of 0 (Table 1).

**Table 1.**
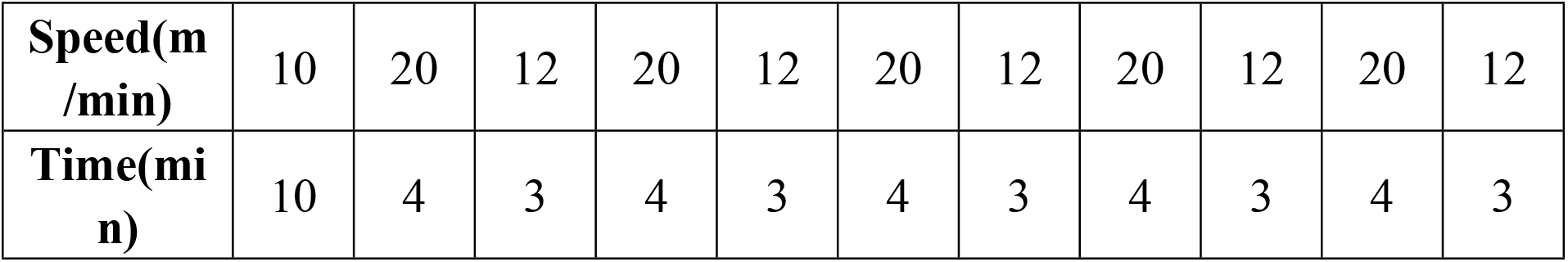
Moderate-Intensity Intermittent Training Protocols.

| Speed(m/min) | 10 | 20 | 12 | 20 | 12 | 20 | 12 | 20 | 12 | 20 | 12 |
| --- | --- | --- | --- | --- | --- | --- | --- | --- | --- | --- | --- |
| Time(min) | 10 | 4 | 3 | 4 | 3 | 4 | 3 | 4 | 3 | 4 | 3 |

### Morris Water Maze Test

The Morris water maze(MWM) experiment had two phases. The first phase was called the “positioning navigation test” and the second was the “spatial exploration test.” These tests evaluated the spatial learning and memory abilities of rats with D-galactose-induced aging^[31]^. The rats were familiarized with the environment one day before the experiment began.For six days, the rats underwent the positioning navigation test. They were placed in the water one at a time from the four quadrants of the maze. Each rat faced the pool wall as it entered. The time it took to find the hidden platform was recorded, with a maximum of 2 minutes allowed. If a rat found the platform, it stayed on it for 10 seconds. If it did not find the platform, the rat was gently guided to it and remained there for 10 seconds before the next trial.On the seventh day, the platform was removed for the spatial exploration test. In this test, the entry point was in the quadrant opposite to where the platform used to be. Over a period of 1 minute, the researchers recorded two things: the percentage of time the rat spent in the target quadrant and the number of times it crossed the previous platform location. The experiment was conducted three times each day until the rats could reliably find the hidden platform.The water maze apparatus was purchased from Anhui Zhenghua Bio-Instrument Co., Ltd.(Suixi, China). It featured a circular pool that was about 0.6 m high and 1.5 m in diameter. The inner walls of the pool were painted black. It also came with a movable, submerged hidden platform. The water temperature was kept at 24±1°C.A blue curtain surrounded the water maze. This curtain blocked external light and created a dim interior. It prevented the rats from using environmental cues to find the platform. A miniature camera (WV-CP500D from Panasonic, Kadoma, Japan) was mounted directly above the pool. The camera connected to a computer and automatically recorded the rats’ movement trajectories.

### ELISA

The prefrontal cortex of the rats were weighed and cut into pieces. They were homogenized in pre-cooled PBS with a protease inhibitor at a ratio of 1:9. Next, the mixture was centrifuged at 12,000 g under 4 °C, and the supernatant was collected. The optical density at 450 nm was measured using an enzyme labeling instrument. This was done following the instructions of the SOD (ZC-36451), GSH-Px (ZC-36648), and MDA (ZC-36429) kits purchased from ZC-36451 Biotechnology Co., Ltd. (hanghai, China) Three samples per group were tested, and each test was repeated three times to reduce error.

### Toluidine Blue Staining

The brain tissue was fixed with 10% neutral formaldehyde and then embedded in paraffin. After routine dewaxing, the tissue was dipped in toluidine blue at 56 °C for 20 minutes. It was lightly washed with distilled water, dipped in 70% ethanol for 1 minute, and then briefly treated with 95% ethanol for rapid differentiation. Next, the tissue was dehydrated using anhydrous ethanol, made transparent with xylene, and sealed with neutral gum. Brain tissue images were collected using a digital slice scanner (Pannoramic 250, 3DHISTECH, Hungary). Each slice was first observed at 4× magnification, and then three pictures at 40× magnification were taken in the prefrontal cortex area.

### Tunel Staining

The fixed brain tissue was first dehydrated with ethanol. It went through 75% ethanol for 4 hours, 85% for 2 hours, 95% for 1 hour, and 100% for 2 hours. Next, the tissue was treated with xylene for 20 minutes. It was then embedded in paraffin for 6 hours. After that, the tissue was dewaxed to water. It underwent citric acid repair for 8 minutes and was cleaned with PBS three times for 5 minutes each. In the dark, a fluorescence Tunel incubation solution was prepared with a ratio of A to B as 1:30. The sample was incubated at 37°C for 1 hour. It was washed with PBS three times (5 minutes per wash). Then, the sample was stained with DAPI for 15 minutes, rinsed with PBS, and sealed with glycerol gelatin. The apoptotic cells in the prefrontal cortex were observed under a fluorescence microscope at 400× magnification and photographed. The apoptosis rate was calculated by dividing the number of apoptotic cells by the total number of cells and then multiplying by 100%.

### miRNA sequencing

Three plasma samples were selected from each group. Total RNA was extracted using the exoRNeasy Serum/Plasma Midi Kit (Qiagen #77044). RNA quality was checked with the Agilent Bioanalyzer 2100 (Agilent Technologies, Santa Clara, CA, US). The RNA was then quantified using the Qubit® 3.0 Fluorometer and NanoDrop One. A small RNA-Seq library of 9 samples was constructed afterward. Cluster formation and first-direction sequencing primer hybridization were completed using the cBot standard process of the Illumina Xten sequencer. The sequencing process used supporting reagents. Data analysis was performed in real time using Illumina’s data acquisition software. The sequencing data were normalized. DESeq2 was then used for data homogenization and analysis of differential expression. Differential miRNAs were identified using the criteria of *P*<0.05 and |log2FC|≥1.

### qRT-PCR

Total RNA was extracted from the prefrontal cortex using the Cell/Tissue Total RNA Kit. We quantified the RNA with a NanoDrop One spectrophotometer from Nanodrop Technologies (Wilmington, DE, USA). We then converted the RNA to cDNA using the Strand cDNA Synthesis Kit. The entire amplification reaction followed the instructions of the SYBR Green qPCR Master Mix. We purchased all the reagents from Yisheng Biotechnology (Shanghai) Co., Ltd. Next, we analyzed the CT values of each sample with Thermo Scientific Piko Real software. We calculated the relative expression of miR-34a based on U6 and the expression of Notch1 mRNA based on GAPDH using the 2^−ΔΔCt^ method. To reduce testing error, we repeated the process three times for three samples from each group. Primer design: Gene sequences were searched in the miRBase database (http://www.mirbase.org) and GenBank. We used the software Primer5 to design the primers. Then, Sangon Biotech (Shanghai) Co., Ltd. synthesized them. The primer sequences are shown in Table 2.

**Table 2.**
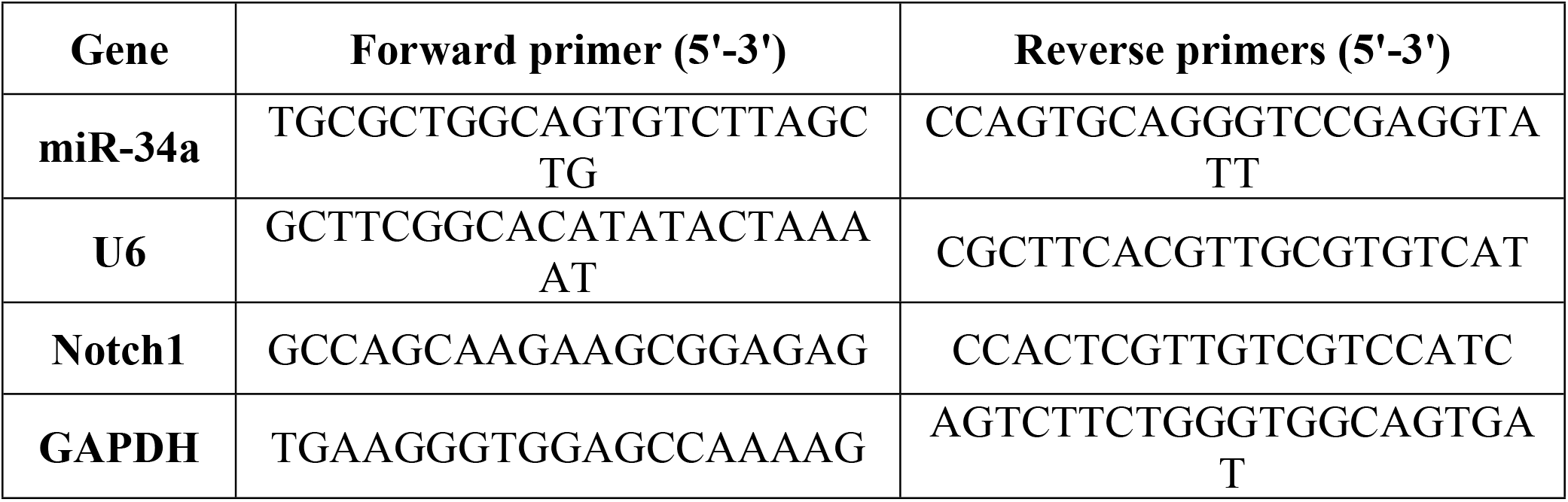
Primer sequences for qRT-PCR.

### Luciferase

We predicted the binding sites for miR-34a and the Notch1 3’UTR using the TargetScan website (https://www.targetscan.org/vert_80/). We then built a Notch1 luciferase plasmid. Next, we co-transfected HEK293T cells with the pcDNA3.1-Notch1 plasmid and either miR-34a mimics or vector mimics using the H4000 reagent. After 48 hours, we lysed the cells. We measured luciferase activity with the Dual Luciferase Reporter Gene Assay Kit from Yisheng Biotechnology Co., Ltd. (Shanghai,China).The relative luciferase activity was calculated by dividing the firefly luciferase activity by the Renilla luciferase activity.

### Western Blotting

We lysed the prefrontal cortex completely using RAPA lysate (P0013C, Beyotime Biotechnology Co., Ltd.(Shanghai, China). Next, we measured the protein concentration with a BCA protein kit (A8020, Beyotime Biotechnology Co., Ltd.(Shanghai, China). We separated the protein using a 10-12% SDS-PAGE gel (PG110, Yamei Biotechnology Co., Ltd.(Shanghai, China). Then, we transferred it to a PVDF membrane by vertical electrophoresis. We blocked the membrane with 5% BSA for 1 hour at room temperature. We added rabbit anti-polyclonal antibodies: Notch1 (10062-2-AP, 1:2000, Proteintech, USA), Bcl-2 (ab194583, 1:500, Abcam), Bax (2772S, 1:1000, CST), and GAPDH (D16H11, 1:5000, CST). Finally, we incubated the membrane overnight at 4°C. The next day, we incubated the anti-rabbit secondary antibody (ZB-2306, 1:5000, Zhongshan Jinqiao Biotechnology Co., Ltd.(Beijing, China) at room temperature for 2 hours. We then washed the membrane with TBST. Next, we developed the ECL luminescent liquid and scanned the gel using a chemiluminescence imager (UVP, BIOSPECTRUM, USA). Finally, we analyzed the gray value of the target band with Image J software and compared the average gray values of the target protein and the GAPDH band between groups.

### Statistical Analysis

All experimental data were entered and collated using Excel 2019. We used GraphPad Prism 9.5 for statistical analysis and to create statistical charts. We compared group differences using one-way analysis of variance (ANOVA). Data are expressed as means ± standard deviation (SD), with statistical significance defined as *P*< 0.05.

## Results

### MIIT can Improve the Spatial Learning and Memory of D-gal-induced Aging Rats

Before MIIT intervention, the escape latency of rats in all three groups generally shortened over time. There was no significant difference between the first and second days. From the third day, groups D and DE had a significantly longer escape latency than group C (*P*<0.01) (Fig 2A). In addition, groups D and DE had fewer platform crossings compared to group C (*P*<0.01) (Fig 2B).After eight weeks of MIIT, a MWM test was repeated to assess the effect of exercise on the rats’ spatial learning and memory. The escape latency in the DE group was significantly shorter than in the D group (*P*<0.05). From the fourth day onward, there was no significant difference between the DE group and the C group (*P*>0.05) (Fig 2D). At the same time, the DE group exhibited more platform crossings than the D group (*P*<0.05) (Fig 2E).

**Fig 2.**
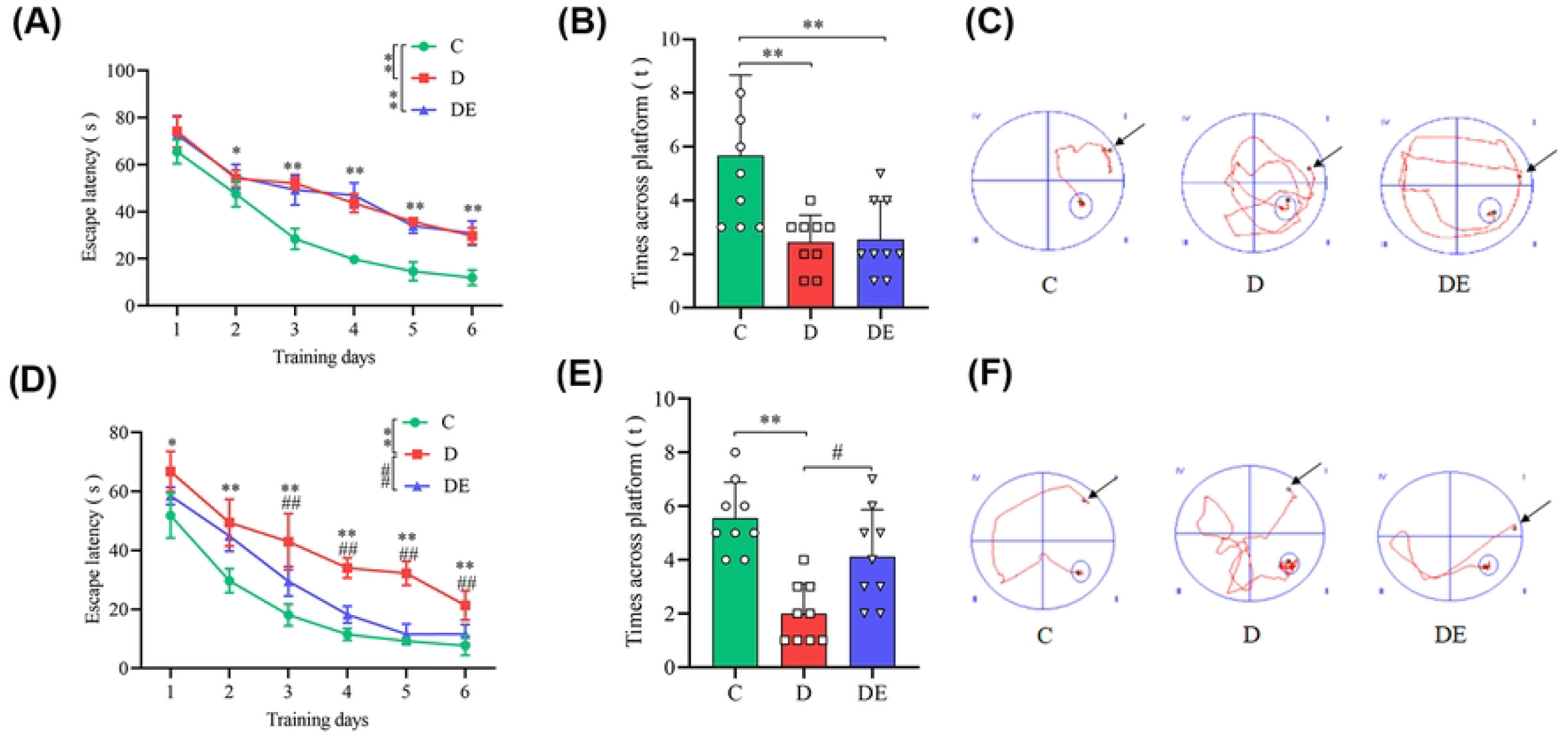
Comparison of MWM tests before and after exercise in each group. (**A**)represents is the average escape latency before exercise. (**B**)indicates is the number of crossingplatform thecrossings platform before exercise. (**C**)shows is the swimming trajectory before exercise. (**D**)represents is the average escape latency after exercise. (**E**)indicates is the number of crossingplatform thecrossings platform after exercise. (**F**)shows is the swimming trajectory after exercise. (mean values±SD, n = 9, ^*^*P*<0.05 vs C, ^**^*P*<0.01 vs C ; ^#^*P*<0.05 vs D ; ^##^*P*<0.01 vs D). C, control group; D, D-galactose group; DE, D-galactose+exercise group.

### MIIT can Increase the Content of SOD and GSH-Px in the PFC of Aging Rats

In group D, the levels of SOD and GSH-Px in the PFC were significantly lower than in group C (*P*<0.01). The MDA level was significantly higher (*P*<0.01), and brain peroxidation products increased. In group DE, the levels of SOD and GSH-Px in the PFC significantly increased compared to group D (*P*<0.01). The MDA level significantly decreased (*P*<0.01) (Fig 3A).

**Fig 1.**
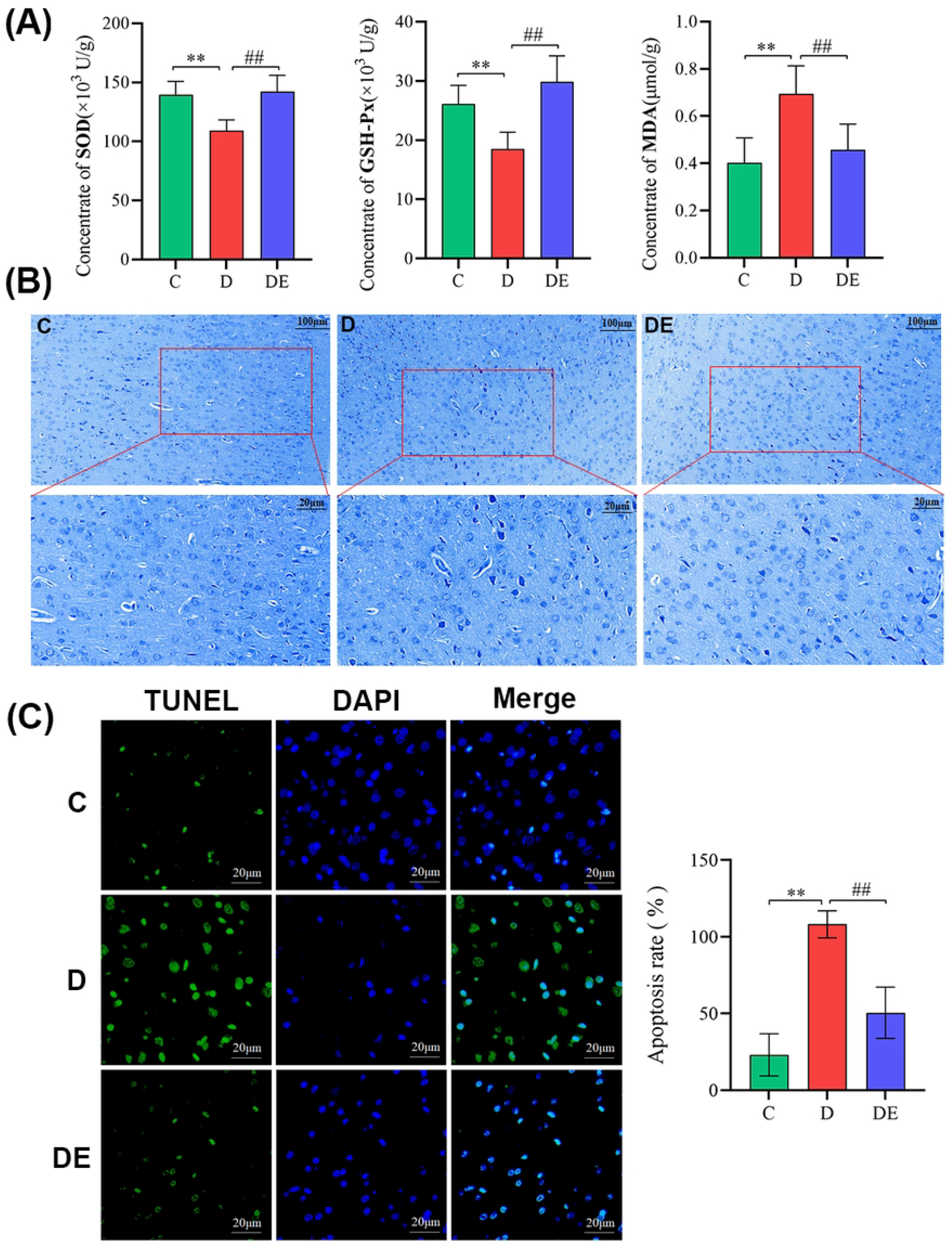
Results of ELISA, Toluidine blue staining, and TUNEL staining in the rat groups. (**A**) shows the levels of MDA, SOD, and GSH-Px in each rat group (U/mg, mean values±SD, n=6, ^**^*P*<0.01 vs C; ^##^*P*<0.01 vs D).(**B**) compares toluidine blue staining in the prefrontal cortex across the groups(40×,scalebar: 20μm). In these samples, the Nissl bodies appear dark blue, while the background is colorless. This happens because the acidic substances in the tissue bind with the cation in toluidine blue, staining the cells blue. Positive expression appears as hypertrophic cells with dark blue round or oval nuclei. (**C**) displays Tunel staining with color fluorescence labeling for apoptosis in the prefrontal cortex cells (40×, scalebar: 20μm). In the images, apoptotic nuclei show green fluorescence, and normal nuclei appear with blue fluorescence (mean values±SD, n=6, ^**^*P*<0.01 vs C; ^##^*P*<0.01 vs D). C, control group; D, D-galactose group; DE, D-galactose+exercise group.

### MIIT can Improve the Morphological Structure of the PFC and Reduce Neuronal Apoptosis in Aging Rats

Toluidine blue staining showed that prefrontal neurons in group C had a complete shape. Their nucleoli were round and obvious. The neurons were neatly arranged and close together, and Nissl particles were large and abundant.In group D, neurons were scattered and sparse. Most had an irregular triangle or polygon shape. Their nucleoli were blurred, and the nuclei were hard to distinguish from nearby tissues. Many cells were vacuolated, and their numbers were significantly reduced. The Nissl bodies also became smaller.In the DE group, the structure of prefrontal neurons appeared more complete. They were arranged in several layers. The nucleoli were clear, and the Nissl bodies were abundant and distinct (Fig 3B).

TUNEL staining showed different patterns. In group D, many apoptotic frontal lobe cells with green fluorescence were observed. The apoptosis rate was much higher than in group C [(23.10±13.76)% vs (108.15±8.80)%, *P*<0.01]. Compared to group D, group DE had noticeably fewer green fluorescent cells, and the apoptosis rate dropped significantly [(108.15±8.80)% vs (50.45±16.66)%, *P*<0.01] (Fig 3C).

### miRNA sequencing was used to screen differential miRNAs in each group of Rats

In the volcano plot, the same miRNA may originate from different precursors. This causes some points to overlap. In group D, compared to group C, 12 miRNAs were differentially expressed. Out of these, 8 were up-regulated and 4 were down-regulated (Fig 4A). In group DE, compared to group D, 14 miRNAs were differentially expressed. Among these, 5 were up-regulated and 9 were down-regulated (Fig 4B). There were more differential expressions between groups D and DE. The heat map shows these specific miRNAs. Blue labels indicate a significant increase in miRNA expression, while red labels indicate a significant decrease.

**Fig 4.**
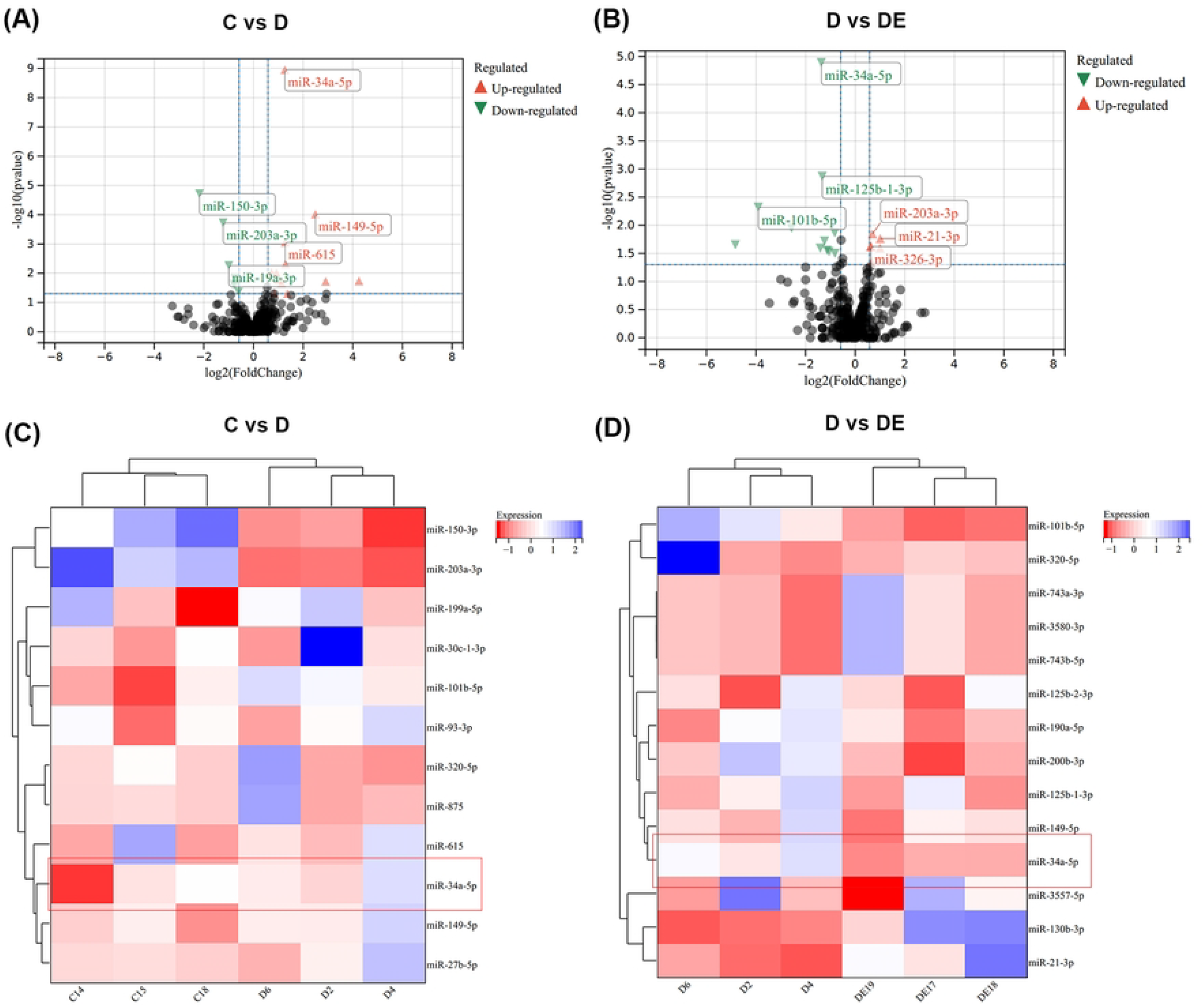
Differential expression of miRNAs in each group of rats. (**A**) and (**B**) display volcano plots of the miRNA expression(The vertical axis, labeled “-log10 *P*-value” represents the corrected *P* value. The horizontal axis, labeled “-Log2 fold change,” shows the expression differences). (**C**) and (**D**) show heat maps of the differential miRNA expression. C, control group; D, D-galactose group; DE, D-galactose+exercise group.

### MIIT can Inhibit miR-34a-5p to Activate Notch-1

First, We predicted a complementary binding between miR-34a-5p and a segment of Notch1’s 3’UTR (positions 190–196) using online bioinformatics tools (Fig 5A). Next, We then measured miR-34a-5p and Notch1 mRNA levels in the rat prefrontal cortex. In group D, miR-34a-5p expression was higher than in group C (1.54±0.57 vs 0.87±0.18, *P*<0.01). Notch1 mRNA expression was lower in group D compared to group C (0.34±0.06 vs 0.94±0.27, *P*<0.01). In the DE group, miR-34a-5p levels dropped compared to group D (1.06±0.28 vs 1.54±0.57, *P*<0.05). At the same time, Notch1 mRNA levels increased (0.70±0.44 vs 0.34±0.06, *P*<0.05) (Fig 5B). Finally, our dual luciferase reporter gene assay confirmed these findings. Co-transfection with miR-34a-5p mimics and the Notch1-PGL4.1 plasmid significantly activated the reporter plasmid’s luciferase activity compared to co-transfection with Notch1 + Vector-PGL4.1 (Fig 5C).

**Fig 5.**
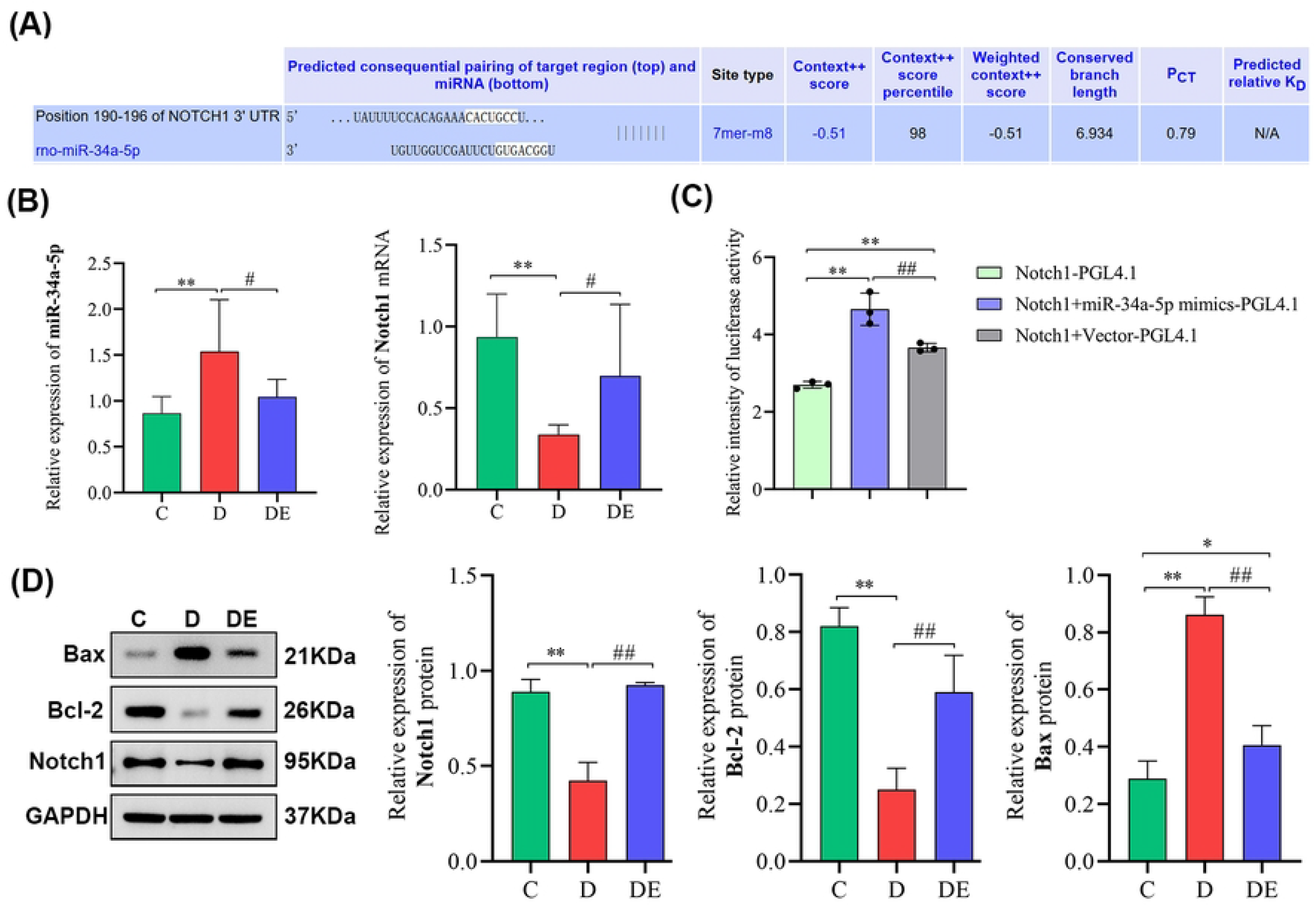
miR-34a-5p and Notch1 binding site prediction, luciferase activity detection and apoptosis-related protein expression level detection results. (**A**) shows the predicted binding sites for miR-34a-5p and Notch1. These predictions were made using the TargetScan website (https://www.targetscan.org/vert_80/). (**B**) shows the levels of miR-34a-5p and Notch1 in the prefrontal cortex of three groups after MIIT (mean values±SD, n=6, ^**^*P*<0.01 vs C ; ^#^*P*<0.05). (**C**) presents the dual luciferase assay results (mean values±SD, n=3, ^**^*P*<0.01 vs C ; ^##^*P*<0.01 vs D). (**D**) shows the expression levels of Notch1, Bcl-2, and Bax in the prefrontal cortex of rats from three different groups (mean values±SD, n=3, ^*^*P*<0.05, ^**^*P*<0.01 vs C ; ^##^*P*<0.01 vs D). C, control group; D, D-galactose group; DE, D-galactose+exercise group.

### MIIT can Reduce the Expression of Apoptosis-related Proteins in the PFC of Aging Rats

In group D, the levels of Notch1 and the anti-apoptotic protein Bcl-2 dropped compared to group C (*P* < 0.01). Meanwhile, the level of the pro-apoptotic protein Bax increased (*P* < 0.01). After 8 weeks of exercise, group DE showed more Notch1 and Bcl-2 than group D (*P* < 0.05). Bax levels were lower in group DE (*P* < 0.05) (Fig 5D).

## Discussion

Aging is an inevitable biological process, and proactive strategies for promoting overall health and maintaining brain vitality have garnered global attention. Accordingly, this study investigates the mechanisms by which exercise delays brain aging in animal models. In the experiment, D-galactose—a compound that simulates the natural process of brain aging—was employed to establish an aging rat model, thereby facilitating the exploration of preventive and therapeutic strategies for aging-related diseases and their underlying molecular mechanisms. The model has been demonstrated to induce oxidative stress, leading to the accumulation of malondialdehyde (MDA), neural damage, and disruptions in the initiation and function of apoptosis-related proteins, thereby impairing cognitive abilities in a manner similar to that observed in natural aging processes^[32]^. In this experiment, administration of D-galactose for six weeks resulted in significant accumulation of MDA in brain tissue compared to the control group, in addition to markedly impairing spatial learning and memory in the Morris water maze test. Furthermore, the observed behavioral changes in the rats were consistent with those reported in previous studies. These findings demonstrate the successful establishment of an aging model in SD rats induced by D-galactose^[33,34]^. Following MIIT, the level of MDA in rat brain tissue significantly decreased, while SOD and GSH-Px levels increased. Additionally, there was a reduction in escape latency and an increase in the number of platform crossings. These findings suggest that MIIT modulates neurochemical profiles, thereby enhancing brain function and improving spatial memory.

MicroRNAs (miRNAs) are non-coding RNAs that bind to target genes and regulate their expression, thereby playing a critical role in a range of biological processes including cell proliferation, differentiation, and apoptosis^[35]^. Alterations in miRNA expression levels in various human diseases further suggest their involvement in the pathogenesis of these conditions^[36,37]^. In this study, we analyzed the expression of miR-34a-5p in brain tissue from each group using quantitative polymerase chain reaction (qPCR). Significant differences were observed between the groups, suggesting that exercise intervention modulates miRNA expression in the brain. The microRNA-34 (miR-34) family comprises three paralogous members: miR-34a, miR-34b, and miR-34c. miR-34a is predominantly expressed in brain tissue, whereas miR-34b/c is primarily found in the lung. Additionally, miR-34a is characterized by multiple subtypes, notably miR-34a-5p and miR-34a-3p, which perform distinct regulatory roles in various tissues and disease contexts. These subtypes show different regulatory roles in different tissues and diseases^[38]^. Recent studies have demonstrated that miR-34a is significantly overexpressed in both the aging process and neurodegenerative diseases related to aging^[39,40]^. Studies have showndemonstrated that the expression of miR-34a-5p in both plasma and brain tissuetissues of is AD upregulated in Alzheimer’s disease patients and several AD mouse models is up-regulated^[41,42]^. We screened for different miRNAs in rat groups using high-throughput sequencing. Our findings showed that MIIT altered the miRNA levels in plasma. Notably, it down-regulated miR-34a-5p. Could lowering miR-34a-5p reduce nerve cell apoptosis? This result motivates us to explore the target genes of miR-34a-5p during exercise interventions that delay brain aging.

At present, the potential target genes related to miR-34a-5p include apoptosis-related genes, such as Bcl-2, Notch and SIRT1, and have attracted much attention because of their involvement in the regulatory mechanism of apoptosis^[43–45]^. These genes are important because they help control apoptosis. Notch1 plays a key role in pathways that lead to cell death. After cerebral ischemia-reperfusion injury, low levels of Notch1 are linked to increased neuronal apoptosis. In contrast, when Notch1 is overexpressed, neuronal apoptosis decreases significantly^[46]^. Research has shown that miR-34a-5p levels rise in rats with myocardial ischemia-reperfusion injury. Inhibitors of miR-34a-5p can target the Notch1 pathway. They reduce the accumulation of reactive oxygen species and inhibit apoptosis. This leads to attenuation of the myocardial ischemia-reperfusion injury^[47]^. We constructed in vitro cell experiments. A dual luciferase assay confirmed that miR-34a-5p targets Notch1 and regulates its activity. High levels of Notch1 increase the anti-apoptotic protein Bcl-2 and decrease the pro-apoptotic protein Bax^[48]^. Peng et al. showed similar results. They found that Notch1 activation raises Bcl-2 levels and lowers Bax levels. This helps recover neurological function and inhibits neuronal apoptosis in depression model rats^[49].^ In our study, the prefrontal cortex of D-gal-induced aging rats showed significantly increased miR-34a-5p. In contrast, Notch1 and Bcl-2 levels declined. These findings suggest that increased miR-34a-5p may worsen apoptosis by inhibiting Notch1. This process might lead to reduced spatial learning and memory in rats.

Regular and appropriate exercise is essential for healthy aging. It reduces the risk of death, chronic diseases, and premature death. Many studies show that exercise helps prevent aging-related diseases^[50–52].^ However, we still need more evidence to fully understand how it works. Research suggests that exercise increases specific miRNAs. These miRNAs travel to target organs and regulate proteins. They may do this by stimulating the body’s response to low oxygen and blood flow. This process helps the body adapt to exercise^[53,54]^. Bao et al. used microarray analysis. They found that 32 miRNAs showed different expression levels in the hippocampus of healthy mice after exercise. Twenty were up-regulated and 12 were down-regulated^[55]^. Exercise may trigger the Notch1 signaling pathway by altering gene expression. This change could improve spatial learning and memory in WT mice and boost synaptic plasticity^[56]^. Other studies show that voluntary wheel running increases miR-130a levels. It also activates autophagy and stops apoptosis. This exercise down-regulates the p53/p21 axis, which helps protect neurons and may slow brain aging^[57]^. In the middle cerebral artery occlusion/reperfusion (MCAO/R) model, exercise reduces neurological dysfunction and infarct size. At 24 hours, it increases the plasma exocrine miR-124 content. Exercise also inhibits STAT3 expression and boosts anti-apoptotic BCL-2 levels. It lowers the levels of the pro-apoptotic BAX protein. As a result, apoptosis is reduced^[58]^. However, the effect of exercise on brain cell apoptosis through miR-34a-5p has not been reported. We conducted MIIT for 8 weeks. This treatment significantly decreased miR-34a-5p expression in the prefrontal cortex of D-gal-induced aging rats. Meanwhile, Notch1 mRNA expression increased. These results indicate that the exercise inhibited the secretion and transcription of miR-34a-5p. To see if these changes benefit the morphological structure of the PFC, we examined the arrangement of prefrontal neurons and measured apoptosis levels. We found that the neurons were neatly arranged. The expression of Notch1 and Bcl-2 proteins increased. Also, the apoptosis rate significantly decreased. MIIT decreases miR-34a-5p expression and reduces neuronal apoptosis. Exercise has a protective effect on the brain. This training may prevent miR-34a-5p from binding to Notch1. It helps regulate the death of prefrontal neurons in aging rats. As a result, spatial learning and memory improve, and brain aging is delayed.

## Conclusion

The study found that eight weeks of MIIT improves spatial learning and memory in aging rats induced by D-gal. It also reduces damage and prevents the death of prefrontal neurons. The likely mechanism involves miR-34a. This molecule targets Notch1 and negatively regulates targets related to cell survival. It helps lower the cell death issues caused by aging and slows down the brain aging process.

## Acknowledgments

We are thankful to all the participants of the study.

## Author Contributions

Conceptualization: Qiaojing Gao, Lu Wang.

Data curation: Jinmei Zhang, Liting Lv, Jun Gao.

Funding acquisition: Lu Wang, Xue Li.

Methodology: Qiaojing Gao, Jinmei Zhang, Liting Lv, Jun Gao.

Validation: Lu Wang, Meng Li, Xue Li.

Writing– original draft: Qiaojing Gao, Meng Li.

Writing– review & editing: Jinmei Zhang, Jun Gao, Xue Li.

